# Spatial structure of natural boxwood and the invasive box tree moth can promote coexistence

**DOI:** 10.1101/2020.11.18.388322

**Authors:** Léo Ledru, Jimmy Garnier, Christiane Gallet, Camille Noûs, Sébastien Ibanez

## Abstract

In the absence of top-down and bottom-up controls, herbivores eventually drive themselves to extinction by exhausting their host plants. Poorly mobile herbivores may experiment only local disappearance, because they can recolonize intact plant patches elsewhere, leaving time to previously over-exploited patches to regrow. However most herbivores such as winged insects are highly mobile, which may prevent the formation of spatial heterogeneity.

We test if long-distance dispersal can preclude coexistence using the invasion of box tree moth (*Cydalima perspectalis*) in Europe as a model system. We build a lattice model and estimate the parameters with a combination of field measurements, experimental data and literature sources. Space corresponds either to a realistic boxwood landscape in the Alps, or to theoretical landscapes of various sizes.

We find that both species persist under a large range of realistic parameter values, despite a severe reduction in boxwood biomass, with an alternation of outbreaks and near-to-extinction moth densities. Large landscapes are necessary for coexistence, allowing the formation of spatial structure. Slow plant regrowth combined with long-distance dispersal could drive moths to extinction, because of resources depletion at the global scale even without a complete synchronization of the local dynamics. The spatial dynamics leads to formation of small plant patches evenly distributed in the landscape, because of a combination of local plant dispersal and global indirect competition between plants through their positive effect on moth population size. Coexistence is favored by such heterogeneous landscapes, because empty patches increase moth mortality during dispersal: the system thus creates its own stability conditions.

## Introduction

In general, most herbivores do not exhaust their resources because they are top-down controlled by their predators [Hairston et al., 1960] as well as bottom-up limited by the defense compounds and the poor nutritional quality of plants [Polis, 1999]. However, in some cases such top-down and bottom-up mechanisms are insufficient to regulate herbivore populations, turning the green world to brown. In such cases, it has been suggested that the spatial dynamics of plant-herbivore metacommunities may favor their coexistence [Wilkinson and Sherratt, 2016]. This hypothesis builds upon long-standing theoretical work which has shown that spatial structure promotes the persistence of otherwise unstable prey-predator systems [Hassell et al., 1991, Comins and Hassell, 1996, Amarasekare, 2008], thanks to local extinctions followed by recolonization, in line with metapopulation and metacommunity dynamics [Hanski and Gilpin, 1997, Holyoak et al., 2005]. These theoretical predictions have received robust empirical support from experiments based on an animal prey-predator system [Taylor, 1991], as protists [Holyoak and Lawler, 1996, Fox et al., 2017] and field studies with arthropods [Nachman, 1988, Winder et al., 2001].

However, there is little evidence showing that the spatial dynamics resulting from interactions between plants and herbivores leads to a green world - or at least to a bi-coloured world with green and brown patches. Many herbivorous insect populations persist thanks to metapopulation dynamics [Tscharntke and Brandl, 2004], but this persistence generally relies on other mechanisms than the depletion of plant resources. For instance, local extinctions can depend on patch size [Eber and Brandl, 1996], on plant resources fluctuations (but for other reasons than the herbivore itself [Halley and Dempster, 1996]), or on a combination of ecological succession and catastrophic events [Stelter et al., 1997]. In the well studied ragwort / cinnabar moth system, the moth can locally disappear following defoliation, but plant patches persist [Myers and Campbell, 1976, Myers, 1976]. Although cinnabar moths contribute to local plant extinction, local plant persistence ultimately depends on habitat suitability, which leads to a source-sink dynamic rather than to a classical metapopulation scenario [van der Meijden and van der Veen-van, 1997, Van der Meijden, 1979]. Moreover, the high dispersal ability of cinnabar moths prevents asynchronous local dynamics for the moth, which rules out a metapopulation model of coexistence [Harrison et al., 1995, van der Meijden and van der Veen-van, 1997]. As far as we know, the only documented plant-herbivore system where the plant goes locally extinct due to over-exploitation comprises the Apiaceae *Aciphylla dieffenbachii* and the monophagous weevil *Hadramphus spinipennis*, two species endemic to the Chatham Istands (New Zealand). Increased local weevil densities are associated with local plant extinction [Schöps, 2002], and numerical simulations have shown that spatial structure allows the persistence of the system, provided that the dispersal distance of the herbivore is intermediate [Johst and Schöps, 2003]. However, the ecological conditions which promote the persistence of this particular system may not hold for other plant-herbivore interactions. In particular, the weevil *H. spinipennis* is wingless and of a large size, which considerably reduces its dispersal ability either by itself, by wind or by birds. In contrast, many insects can disperse over long distances [Wilson and Thomas, 2002, Gillespie et al., 2012]. Long-distance dispersal can promote metapopulation persistence, except when strong density dependence triggers local extinctions [Johst et al., 2002]. In that case, long-distance dispersal events synchronize local extinctions which eventually lead to the extinction of the whole metapopulation [Palmqvist and Lundberg, 1998, Johst et al., 2002]. In plant-herbivore metacommunities, strong density dependence occurs when herbivores over-exploit their host down to local extinction.

In order to test if plant-herbivore metacommunities can persist despite high abilities of herbivores to disperse, we study the system formed by the common European boxwood *Buxus sempervirens L*. and the invasive box tree moth *Cydalima perspectalis* (Walker, 1859)(Lepidoptera: Crambidae) in Europe. This invasive species first arrived and established in Germany in 2006/2007 [Van der Straten and Muus, 2010] human assisted via the boxwood trade from Asia [Kenis et al., 2013, Van der Straten and Muus, 2010, Bras et al., 2019]. Then, the moth rapidly spread over Europe [Blackburn et al., 2011], invading almost the entire European buxus range by 2021 as expected [Nacambo et al., 2014]. This rapid invasion suggests the occurrence of human-mediated and natural long-distance dispersal events. Moreover, this species causes severe defoliation mainly because its fecundity is high and the individuals lay masses of eggs directly on the leaves. Thus, defoliation caused by moth larvae can lead to the death of the boxwood, especially when the bark is also consumed [Kenis et al., 2013]. After this possibly total defoliation, boxwood can either grow back or wither completely and die if the defoliation becomes too recurrent or when bark is consumed [Kenis et al., 2013]. This harmful impact of the moth may have major economic consequences at long terms [Mitchell et al., 2018]. The local extinction of boxwood has already been observed in the Nature Reserve of Grenzach-Whylen in Germany [Kenis et al., 2013]. In the meantime, the moth goes extinct locally after total defoliation of boxwood stands, even if it grows back several years after the moth outbreak. Within its home range, the moth, which also consumes boxwood, is regulated by its natural enemies [Wan et al., 2014] and no local extinction of either plant and insect species is observed, but potential European natural enemies do not significantly alter the invasive moth population dynamics [Kenis et al., 2013, Leuthardt and Baur, 2013]. Moreover, although box trees contain highly toxic alkaloids [Ahmed et al., 1988, Loru et al., 2000, Devkota et al., 2008], the moth larvae can sequester them in their body [Leuthardt et al., 2013]. Thus, in absence of top-down and bottom-up control of this invasive species, the question remains: will the European boxwood stands remain green? In contrast to our system, the question does not arise with the ragwort / cinnabar and Apiaceae / weevil systems mentioned earlier since both insects are native and may therefore be top-down controlled by local natural enemies. However, in North America, Australia and New Zealand, the ragwort and the cinnabar are both non-native, thus they are both free of top-down and bottom-up controls which is also different from our system [CABI, 2021].

Metacommunity dynamics with local moth extinctions followed by recolonization may be an alternative mechanism to top-down and bottom-up control favouring coexistence in Europe. In the particular context of biological invasions, spatial effects have not been widely addressed [Melbourne et al., 2007], although they may favour coexistence. The metacommunity mechanism requires spatial heterogeneity among local communities, which is likely because of the fairly fragmented distribution of boxwood in Europe, and because of the temporal shift between different patch invasion. As long as the invasion does not start simultaneously in all stands, the moth may disperse from its current totally defoliated stand to a green intact stand. The defoliated stands may then recover and be recolonized lately. Thus, despite local extinctions and recolonizations, local fluctuations may be averaged on a large spatial scale, leading to a global stationary regime called ‘statistical stability’ [De Roos et al., 1991, Holyoak et al., 2005, Amarasekare, 2008]. However, unlike the wingless weevil *H. spinipennis*, the highly mobile *C. perspectalis* can fly or be transported by exogenous factors (wind, human activities) [Bras et al., 2019]. Its high mobility may prevent spatial heterogeneity and therefore precludes coexistence by spatial effects [Johst et al., 2002, Johst and Schöps, 2003]. Thus at large spatial scale, three ecological scenarios are likely to occur. First, the moth might very quickly overexploit its host, causing its own extinction but not the one of its host, which in turn slowly grows back. Second, the moth might persist long enough to exhaust its host, leading to the disappearance of both species. Third, coexistence might result from the balance between local moth extinctions and recolonizations, without complete resource depletion. Our study focuses on the conditions that favor such coexistence, based on the following hypotheses:

1. Long-term coexistence of boxwood and moth is possible at the landscape scale through spatial stabilizing effects (1a). Those effects rely on asynchronous local dynamics (1b).
2. Despite cycles of local extinctions and recolonizations, the coexistence regime is stationary at the regional scale, which corresponds to statistical stability.
3. Dispersal is double-edged: very limited dispersal might prevent the colonization of green patches (3a), whereas long-distance dispersal may synchronize local dynamics (3b).
4. The coexistence regime depends on the landscape properties, in particular the landscape size and the proportion of boxwood patches in the landscape. First, larger landscapes favor coexistence (4a). Secondly, the effect of the proportion of boxwood patches is uncertain, since it provides more resources to the moth, but also favors outbreaks and resource depletion (4b).

In order to address these four hypotheses, we develop a population model dynamics for the boxwood and moth system. First, a local model reproduces the local invasion dynamics, which invariably leads to moth extinction in the field. Then, a spatially explicit model simulates the dynamics of the moth in a landscape. Our model is calibrated from the literature, *in situ* measures, and through mesocosm experimentation.

## Study system & theoretical model description

### Species involved: boxwood and box tree moth

The box tree moth, *Cydalima perspectalis*, is an herbivorous lepidoptera belonging to the Crambidae family [Mally and Nuss, 2010]. In Europe, five to seven larval instars are necessary to the larvae to become pupae for about ten days before emerging as moths [Kawazu et al., 2010]. During the winter, the larvae are at the beginning of their development in stages two, three or even four [Poitou et al., 2020] and form cocoons to enter in hybernaculi [Nacambo et al., 2014]. The moths live up to two weeks during which they reproduce and lay eggs on boxwood leaves. The moth has a high fecundity rate, with between 300 and 400 eggs laid per female in its native range, and until 800 eggs in Europe in masses of 5 to 30 eggs [Kawazu et al., 2010, Wan et al., 2014, Tabone et al., 2015]. In Asia, two to five life cycles generations are possible per year, with a break during the winter when the caterpillars are dormant [Maruyama et al., 1987, 1991]. In its invasion range in Europe, the moth completes from 2 (in the north) to 4 (in the south) generations per year [Nacambo et al., 2014, Göttig, 2017]. The mean intrinsic dispersal distance of moths has been estimated around ten kilometers per year [Van der Straten and Muus, 2010].

The moth exhibits no preference for any particular boxwood species [Leuthardt and Baur, 2013], so the common European boxwood *Buxus sempervirens* is widely consumed, as well as Caucasus boxwood *Buxus colchica*, and even the rarer European species *Buxus balearica* [Kenis et al., 2013]. These natural boxwood stands, which have already undergone a major decline over the last millennia [Di Domenico et al., 2012], are now subject to this additional threat. In Asia, *C.perspectalis* also consumes other species, including holly (*Ilex purpurea*), charcoal (*Euonymus japonicus*, and *E.alatus*). Fortunately these species do not seem to be consumed in Europe [Göttig, 2017]. Despite natural regulation by native predators and parasites, this moth remains a threat to ornamental boxwood in Asia, where its potential targets are protected by insecticides [Wan et al., 2014]. Instead, in Europe *Bacillus thuringiensis* is commonly used as a sustainable control method. However, its efficiency is offset by its low persistence and more importantly its wide target range. Thus, current efforts are being made to develop more long-term specific treatments. Biological control solutions are also being explored, such as the use of nematodes [Göttig and Herz, 2018] and parasites from the genus *Trichogramma* [Göttig and Herz, 2016]. Efforts are also being made to seek out predators and parasites from the box tree moth’s area of origin that might act in areas of invasion [Göttig, 2017]. The use of pheromone traps is widespread, both for monitoring and control [Santi et al., 2015, Göttig and Herz, 2017], but their effectiveness appears to be insufficient at a large scale. Even if effective control for ornamental boxwood could be introduced, natural boxwood and associated ecosystems will likely suffer dramatically from the *C.perspectalis* invasion.

Boxwood has a fairly heterogeneous distribution in Europe that consists mainly of small and fragmented stands, but some large areas of continuous boxwood occur in the French Pyrenees, the Pre-Alps and the Jura [Di Domenico et al., 2012]. It is a long-lived, slow-growing shrub that thrives on calcareous substrate. It can tolerate a wide gradient of light incidence, and can therefore be found in a range of plant communities, from canopy areas in heaths to under cover in forests [Di Domenico et al., 2012]. It can play an important role in structuring ecosystems, by trapping sediment and storing water. It also influences the establishment and survival of tree species in woodland succession [Mitchell et al., 2018]. A total of 286 species are associated with the shrub, including 43 fungi that are exclusively observed on boxwood [Mitchell et al., 2018]. However, boxwood is scarcely predated by native species.

### Local demographic model

We start with a local model that describes the interaction between BTM and its host BT at a small homogeneous spatial scale. The model projects the population size *m* of Box Tree Moths (BTM) and the population density of Box Trees (BT), which are separated in two variables: leaf density *l* and wood density *w*, from moth generation *n* to *n* + 1. This time representation is used to avoid the problem of multiple generations per year and its spatial variation. However, we are able to project the population of BTM and BT from year to year if we know the number of generations per year in each specific location. We write

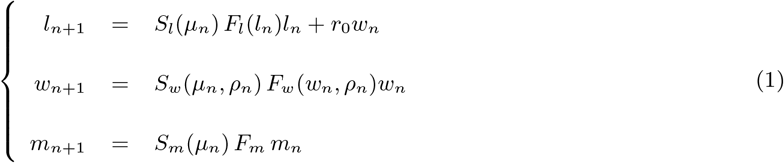

to indicate that during the projection interval, BT and BTM grow and reproduce (F), and survive (S). The BT reproduction functions were constructed using a Ricker model, which includes the intrinsic population growth rates *r_f_, r_w_* and the carrying capacities of the environment *L_max_, W_max_*, while the BTM reproduction function is linear and only includes the fecundity of adults *f* and their survival *s*. The survival of the species is determined by the consumption of leaves and bark by the BTM as well as the intraspecific competition for the resource faced by BTM. The survival and reproduction functions *F* and *S* depend on both the current population *l,w,m*, and the environmental descriptors *μ* and *ρ*.

#### Environmental descriptors

The environmental quality is described using the following two quantities:

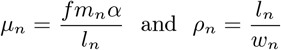

The descriptor *μ* corresponds to the ratio between the number of leaves needed by all the larvae to fulfil their cycle complete development and the number of available leaves (*α* is the amount of leaves needed per larva). The number of larvae depends on the number of moths (*m_n_*) through its product by the moth fecundity (*f*). And each larva needs *α* leaves to complete its cycle. The ratio *μ* thus quantifies the pressure for the resource, which plays a direct role in the intensity of consumption of leaves and wood, and therefore the survival of the larvae.

The descriptor *ρ* is the quantity of leaves per unit of wood. This represents the level of boxwood defoliation, which has an impact on the growth of the wood.

#### Reproduction

The increase in foliage biomass is the result of two processes: the growth of leaves *F_l_*, which depends on the current foliage (Figure 1a), and the production *r_0_* of new shoots by the wood after defoliation (Figure 1b). Without herbivory, the growth of leaves is limited only by senescence and its carrying capacity *L_max_*. The leaves growth *F_l_* is represented by a Ricker model of the following form

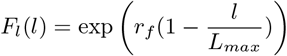

where *r_f_* is the intrinsic growth rate of the leaves.

**Fig. 1.**
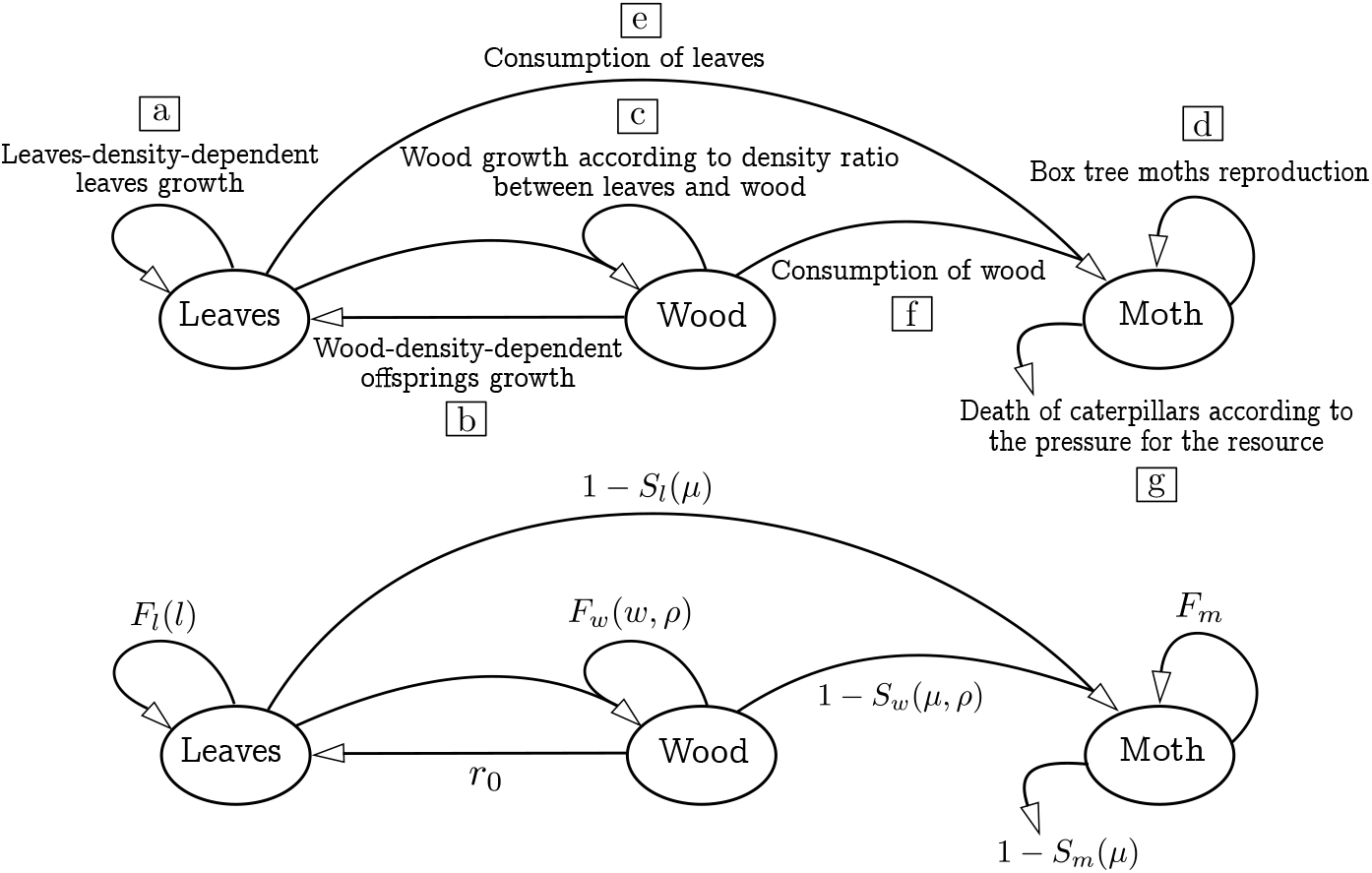
Model of dynamics between the boxwood, separated into wood and leaves, and the box tree moth. The arrows show the interaction between the three variables

The wood growth function *F_w_* (Figure 1c) is constructed using a Ricker model. Positive growth is constrained by carrying capacity. Negative growth occurs after an important defoliation because branches or even a proportion of trunk can die after defoliation. For each projection interval, the intrinsic growth rate of the wood is defined as the balance between the production of new wood, which critically depends on the density of leaves per unit of wood *ρ*, and the mortality induced by severe defoliation. When the density of leaves is large (*ρ* ≫ 1), the BT is healthy and its production of wood reaches a maximum. Conversely, when the density of leaves per unit of wood collapses due to severe defoliation, the production of wood is low while the mortality increases until a maximum which forces the growth rate to be negative (Supporting Information (SI) for details).

The reproduction rate of BTM (Figure 1d) does not suffer from density dependence and results from the product of adult fecundity *f* and adult survival *s*

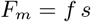

#### Survival

The leaves may die by senescence at rate *v*, or be consumed by BTM at a rate which increases with the pressure of BTM on BT (*μ*) (Figure 1e). In the absence of BTM, 0% of leaves are consumed, while if BTM have saturated the environment, 100% of the leaves are consumed (see SI for details).

The wood can suffer from both defoliation (Figure 1c), which decreases its intrinsic growth rate, and bark consumption by the BTM (Figure 1f). The wood mortality due to consumption increases with both the BTM pressure, *μ* and the BT health *ρ*. More precisely, the wood mortality saturates to *d_max_* when the foliage is abundant. However, when the foliage is small, the bark consumption occurs while the superficial wood is available. Thus recently defoliated boxwood with small bark coverage (*ρ* close to 0) cannot be consumed by the larvae (see SI for details). We use a step function which allows to take into account that the consumption of superficial wood depends on the presence of available softwood, and thus a certain amount of foliage.

Survival of BTM during the larval stage depends mainly on the amount of available resource per larva *μ*. If the larva has enough available resource to complete its six stages, it will evolve into a moth, while a lack of resource during its growth will cause its death. The survival rate also takes into account intraspecific competition for resource caused by interference between the larvae (see SI for details).

### Spatially explicit model

Using the local model which describes the interactions between BTM and BT at small homogeneous spatial scale, relevant for short time dynamics, we build a cellular automaton in order to investigate the BTM invasion over a regional heterogeneous landscape and long time scale. Indeed, while the local model evaluates the moth invasion on the scale of a few generations, the results with explicit space are obtained with a projection time of 1000 generations, i.e. 500 years in the case of two generations per year.

#### Landscape: French Alps and theoretical grid space

Our spatially explicit model describes the dynamics of the BT and BTM on a 2 dimensional landscape composed of two types of cells: habitat cells where BT can grow and urban cells where BT cannot establish. In each cell with BT, BTM can mate and lay eggs according to the local demographic model, while in cells without BT, BTM cannot become established. After the reproduction phase, BTM moths disperse over the landscape to find new areas to breed and lay eggs. BT can also disperse over the landscape to recolonise extinct areas. Simulations are initialized with a single moth invading a patch chosen at random, and carried out on a maximum of 1000 iterations if no specifications are given. We consider two types of landscape: a schematic map of the French-Alps and a theoretical landscape.

The French-Alps map was obtained from field observations provided by the National Alpine Botanical Conservatory. It is composed of 570 by 351 cells of 29 hectares each (about 58 000km^2^, SI Figure S.1). It should be noted that these data focus on natural boxwood and neglect the presence of ornamental boxwood in urban areas. We focus on the French Alps because detailed botanical data are available, but in theory our model could be extended to the whole area of invasion. This landscape allows us to investigate the potential impact of the BTM in this region.

We also consider a more theoretical squared landscape to investigate the effect of the spatial structure on the invasion of the BTM. In particular, we use various initial size and proportion of boxwood patches. For each landscape, we also calculate an aggregation index by counting the number of pairs (adjacent boxwood patches) in the landscape and dividing it by the maximum number of possible pairs: 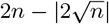, where *n* is the proportion of boxwood cells [Harary and Harborth, 1976]. For each landscape size and boxwood proportion, we randomly generate 1000 landscapes with possibly different aggregation indices. The final aggregation index equals the difference between the index of the landscape of interest and the average index of the randomly generated landscapes.

#### Dispersal phase of BTM

A BTM dispersal event includes two stages: (1) emigrating from birth areas, and (2) searching for new areas (exploration) and settling to breed. Field observations suggest that the exploration phase is stochastic, composed of frequent short-distance dispersal by adult flight, and rare long-distance dispersal by anthropogenic action (boxwood trade) or long flight. *In situ* experimentation using a flight carousel has provided a mean dispersal distance per individual of 13km (Bras et al., personal communication), which is in accordance with the 10km dispersal distance observed by Van der Straten and Muus [2010]. Bras et al. have also observed rare long-distance flights in their experiments, which may correspond to long-distance dispersal events that we model using a fat-tailed dispersal kernel. To this end, we use an exponential power distribution from Klein et al. [2006] which makes it possible to compare different shape of distribution tails while maintaining a fixed average dispersal distance. For ecologically realistic calibration the tail shape parameter is 0.5 (i.e fat-tailed dispersal kernel) and the average dispersal distance is 25 cells (i.e ≈ 13km). In addition, to save computation time, we assume that BTM disperses as a swarm of 1000 individuals. This group dispersal may occur because BTM can be attracted by volatile boxwood compounds or avoid geographical barriers, or are influenced by weather conditions. Thus, during the searching and settling phase, each group of BTM settles in an area located at a random distance drawn in the exponential power distribution and chosen with a random turning angle run in a uniform distribution over (0, 2*π*). The emigration rate for each location depends on the pressure for resource *μ* at the birth location. As long as pressure remains low, the moths have the possibility to find leaves to oviposit in their birth patch and thus dispersal is weak. When the resource pressure increases, there is not enough boxwood available for laying eggs and adults will disperse massively to another patch in search for resource. Such resource-dependent dispersal has also been modelled by the study of Johst and Schöps [2003] (see SI for details).

#### Dispersal of BT

Dispersal events for BT include (1) creation of seeds and (2) dispersal of seeds to surrounding areas by wind and birds. We assume that BT dispersal is very low and occurs only if the boxwood is in fairly good condition, meaning that it has sufficient foliage (see SI for details).

As such, the initial density of wood in a newly dispersed seedling depends on the parent patch density. Foliage is produced in the next generation after recolonization through the production of leaves by the wood. We assume that seeds from a location are transported randomly only in the 8 surrounding cells that has previously contained BT. We only make it possible for an extinct area to be recolonized by surrounding BT, and we make it impossible for BT to colonize new areas.

### Parameter estimation and sensitivity analysis

To quantify the parameters of our model to our system study (Box Tree and Box Tree Moth), we use three different methods: 1) literature review, 2) experimental measurement (field data and mesocosm experiment) and 3) statistical analysis.

#### Literature review

From the recent literature on the Box Tree Moth [Kawazu et al., 2010, Wan et al., 2014, Tabone et al., 2015], we can evaluate the fecundity parameters *f* with a mean value of 120 and a range from 0 (i.e effects of oophagus predators and parasites) to 300 (see SI for more details). In addition, the maximum survival rate of the caterpillar is on average 49%. Then combining experimental results from T. Defferier and E. Tabone from the National Research Institute for Agriculture, Food and Environment (INRAE) and Slansky Jr and Scriber [1982], we estimate the parameter *α* around 0.3 (see SI for more details). From the experimental study of Bras [2015], we are able to quantify the average dispersal distance of the BTM *α_d_* and the tail shape parameter *c*.

#### Experimental results

To assess some parameters describing the interactions between the two species we perform *in situ* measurements and a mesocosm experiment. First, we collect data in two boxwood areas located on the eastern slope of the Epine massif in Savoie (45°38’23.7″N 5°50’43.6″E and 45°41’33.2″N 5°50’56.2″E) at an altitude of 500 and 630 meters, respectively. IN this are, the invasion peak resulting in massive defoliation of the boxwood, occurred in July and August 2016, while we sampled in March 2017. From these measurements on already defoliated area, we estimate from 101 boxwoods its mortality due to leaves consumption by BTM *d_max_* = 0.74 and from 49 boxwoods its shoot production parameter *r*_0_ = 5.10^−5^ (see SI for more details on the parameter estimates). Moreover, from dendrometry method and statistical analysis, we estimate the maximum growth rate of the wood to *r_w,max_* = 0.3. Secondly, we perform a mescosm experiment to model the consumption of leaves by caterpillars and the caterpillar survival with respect to the descriptor parameter *μ*. We created a gradient for the resource pressure *μ* by placing varying numbers of caterpillars on the box trees. The gradient have seven *μ* values (see SI for more details on the experiments and the statistical analysis).

#### Statistical analysis from the local model

In order to quantify the other parameters of the model, we use our local demographic model. Field observations show that when the box tree moth colonizes a patch, its population explodes within a few generations, eventually reaching a peak of density resulting in total defoliation of the boxwood stand. Hence, no more resources are available and the box tree moth disappears from this patch. This dynamics lasts for 3 to 4 years after the beginning of the invasion until the extinction of the moth. When two generations are carried out per year, it corresponds to 6 to 8 generations. Using statistical analysis, we look for parameters that reproduces this qualitative behaviour, and achieves plausible quantitative outputs (see parameters referred as “estimate” in Table 1 and details in SI section S.1.2).

**Table 1:** Model parameters. The values correspond to those most representative of actual ecological conditions. The parameters are either measured quantitatively, i.e. a direct value of the parameter concerned is measured, or qualitatively, i.e. the measurement of a process allows the calibration of the associated parameter. We estimate the unmeasured parameters from the model simulations, and aim to be consistent with the ecological situation. The source listed as “experiment” corresponds to the measurements of moth weights made by the INRAE for *α*, and the mesocosm experiment for *σ_m_, σ_l_* and *S_m,max_*. “Flight carousel experiment” corresponds to the measurements made by Bras et al. from the INRAE.

|  | Settings | Measured | Values | Unit | Meaning | Source |
| --- | --- | --- | --- | --- | --- | --- |
| Leaves | $v$ | No | 0.98 | $\frac{1}{t}$ | Leaf survival rate at senescence | estimate |
| | $r_f$ | No | 0.4 | $\frac{1}{t}$ | Leaf growth rate | estimate |
| | $L_{max}$ | No | $\frac{1}{3}W_{max}$ | $g.ha^{-1}$ | Leaf carrying capacity | estimate |
| | $r_0$ | Quantitative | $5 * 10^{-3}$ | $\frac{1}{t}$ | Offshoot production rate | field |
| | $\sigma_l$ | Qualitative | 0.01 | - | Consumption function setting | experiment |
| Wood | $W_{max}$ | No | $3 * 10^9$ | $g.ha^{-1}$ | Wood carrying capacity | estimate |
| | $d_{max}$ | Quantitative | 0.74 | - | Saturation value of the maximum mortality function $D_{max}(\rho)$ | field |
| | $\beta_s$ | No | 10 | - | Setting of the wood mortality function by consumption (curvature) | estimate |
| | $\theta_s$ | No | 1.2 | - | Setting of the wood mortality function by consumption (inflection point) | estimate |
| | $\beta_r$ | No | 5 | - | Setting of the wood growth function (curvature) | estimate |
| | $\theta_r$ | No | $5 * 10^{-5}$ | - | Setting of the wood growth function (inflection point) | estimate |
| | $r_{w,max}$ | Qualitative | 0.3 | - | Saturation value of the wood growth function | field |
| | $r_{w,min}$ | No | 1 | - | Saturation value of the growth deficit function $\gamma_0$ | estimate |
| | $d$ | No | 0.95 | - | Setting of the growth deficit function $\gamma_0$ | estimate |
| | $\omega_1$ | No | 0.1 | - | Setting of the wood dispersal function | estimate |
| | $\omega_{max}$ | No | $1 * 10^{-5}$ | - | Maximum proportion of wood that can contribute to dispersal | estimate |
| Box tree moth | f | Quantitative | 120 | individual | Box tree moth fecundity | literature<br>[Kawazu et al., 2010]<br>[Wan et al., 2014] |
| | $\alpha$ | Quantitative | 0.3 | g | Amount of leaf needed by a box tree moth for its larval cycle | literature<br>[Slansky Jr and Scriber, 1982]<br>and experiment |
| | $\sigma_m$ | Qualitative | 0.85 | - | Setting of caterpillar survival function | experiment |
| | $S_{m,max}$ | Quantitative | 0.49 | $\frac{1}{t}$ | Maximum caterpillar survival rate | literature<br>[Kawazu et al., 2010]<br>and experiment |
| | $s$ | No | 0.4 | $\frac{1}{t}$ | Survival rate of adults (moths) | estimate |
| | $M_{m,max}$ | No | 0.01 | $\frac{1}{t}$ | Survival rate of adults during dispersal | estimate |
| | $\delta$ | No | 0.01 | - | Parameter of the dispersal function | estimate |
| | $c$ | Qualitative | 0.5 | - | Tail shape parameter of the Exponential power distribution | flight carousel<br>experiment |
| | $\alpha_d$ | Quantitative | 25 | cells | Average dispersal distance of the Exponential power distribution | flight carousel<br>experiment |

#### Sensitivity analysis

First we discuss the outcome of the local demographic model. Although the moth population collapses for realistic parameter values as expected from the observations (Figure S.1 in SI), the model may reach a coexistence equilibrium for some parameter values. In particular, we look at two key parameters, the moth fecundity and moth survival that may vary during the course of the outbreak in Europe. Indeed, the accommodation of native predators and parasites could reduce the survival rate of caterpillars, and moth fecundity could be reduced by oophagous insects. From our local demographic model, we show that coexistence is only possible in a very narrow range of parameters values, which are far from the actual measured values (see Figure S.2 in SI). To go further in the understanding of the coexistence behavior, we look at two connected patches which may favour coexistence in plant-insect interactions [Kang and Armbruster, 2011]. However, our numerical simulations show only a slight expansion of the coexistence area, which is far from sufficient to lead to coexistence with realistic parameters (see Figure S.2 in SI). Thus we really need a spatial model to understand the invasion of the BTM.

To further investigate the effect of BT and BTM interactions and the robustness of the spatially explicit model, we conduct a global sensitivity analysis of three outcomes of our model: the boxwood biomass, the percentage of dead boxwood cells and the probability of moth persistence. For each outcomes, we use the Partial Rank Correlation Coefficients (PRCC) [Saltelli et al., 2004] to quantify the strength of sensitivity for each of the 21 parameters and the direction of their impact on the model outcomes. If the PRCC is positive, the parameter increases the outcome, while it decreases for negative PRCC. We sample parameters in a wide range centered on the reference values of Table 1, that remains ecologically coherent (e.g. boxwood growth is slow). We use 1786 values sampled around the reference values of Table 1 for each parameter and 10 replicates per parameter values to compute the PRCC (see section S.4 and Figure S.9 for more details and global sensitivity results).

### Modelling results and discussion

#### Spatial structure promotes coexistence while resource depletion drives to extinction (Hyp 1)

Using a spatially explicit model, including local population dynamics and short to long range dispersal events, we show that coexistence of the moth/boxwood system occurs across a wide range of parameters. At a regional scale, dispersal allows box tree moth persistence in a cycle outbreak dynamic [Berryman, 1987], through recurrent recolonization of patches that have been previously defoliated and which have had time to recover. The spatial structure therefore allows coexistence, in line with hypothesis 1a. The coexistence mechanism is similar to the rock-paper-scissors game with the corresponding states: patches of defoliated box tree, patches of box tree with foliage and patches of box tree invaded by the moth. These three states compete with each other following a circular hierarchy, as defoliated box trees ‘lose’ against box trees with foliage, which are in turn invaded by box tree moths, which finally leads to defoliated box trees. Similar rock-paper-scissors games have been described in other ecological contexts such as polymorphic bacterial strains [Kerr et al., 2002] and plant-mutualist-exploiter systems [Szilágyi et al., 2009].

The sensitivity analysis (SI Figure S.9) reveals that fecundity and survival of the box tree moth significantly reduce its persistence over the landscape. Thus, we explore moth persistence over a larger range of fecundity and survival parameters than those estimated. Predation on the moth and on the caterpillars is currently low [Kenis et al., 2013], in part because the box tree moth accumulates boxwood alkaloids in its body [Leuthardt and Baur, 2013]. However, native predators may become efficient to feed on the moth following phenotypic plasticity or adaptation [Carlsson et al., 2009]. Native egg parasites like trichograms often used in biological control may also become able to feed on the moth, thereby reducing its fecundity. We find that the moth could rapidly go extinct only for very low fecundity and survival rates (lower-left corners of Figure 2a and f). It is therefore unlikely that the accommodation of native natural enemies will trigger moth extinction. Elevation can also have an important influence on larvae survival and fecundity (via egg survival), as well as on the nutritional quality of the boxwood that controls larvae growth. However, we show that elevation does not change qualitatively the outcomes of the model even if it reduces the above mentioned parameters by 50% (SI Figure S.9). Lastly, the sensitivity analysis shows that moth persistence is significantly impacted by the boxwood productivity and survival (SI Figure S.9). Thus environmental changes induced by human exploitation, climate changes or boxwood parasites, should have a critical impact on the BTM persistence through its effect on boxwood at local and global scale.

**Fig. 2.**
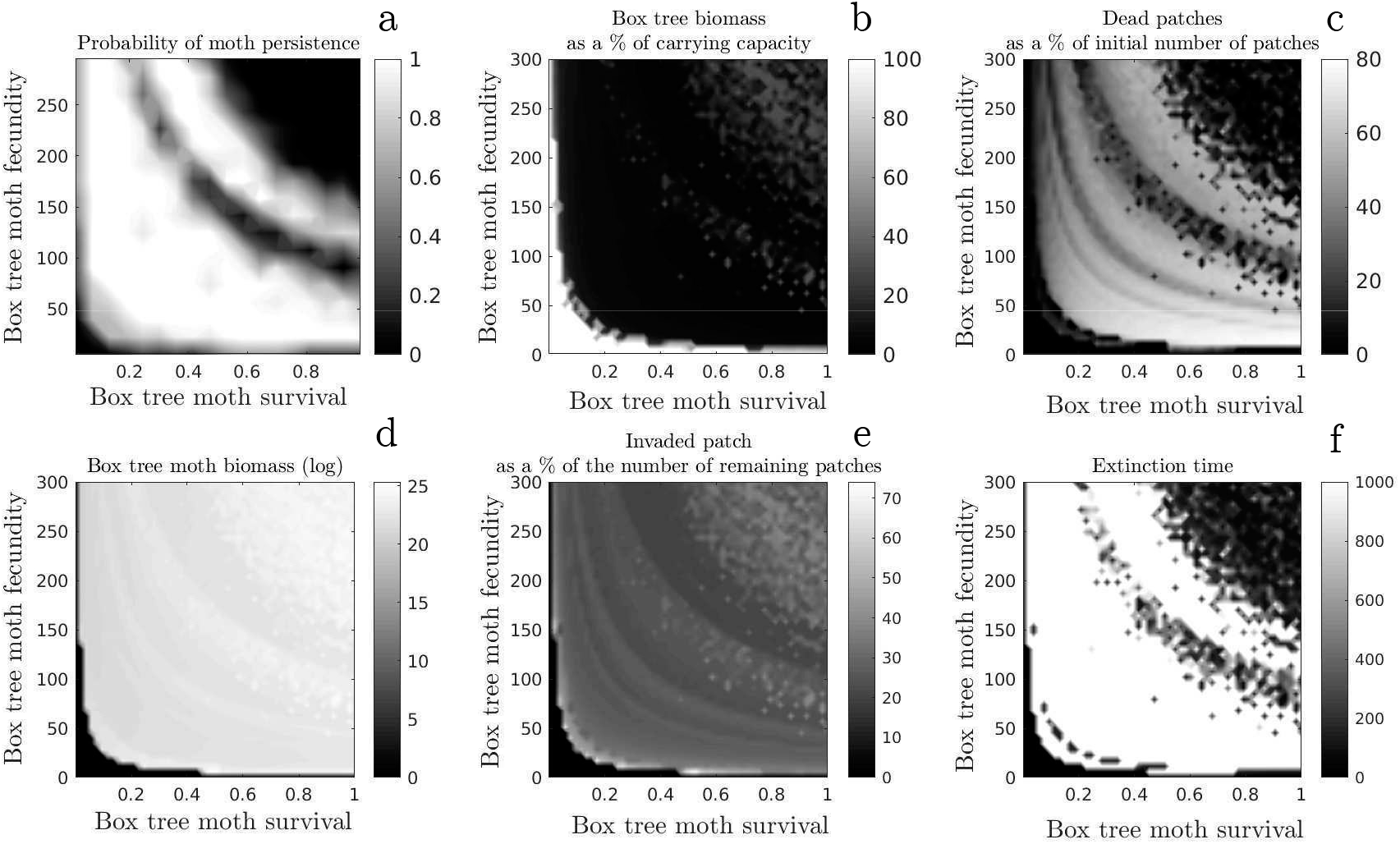
Final state maps in the real landscape in function of fertility *f* and maximum survival *S_m,max_*. (a) landscape-scale probability of moth persistence, (b) landscape-scale wood biomass expressed as a percentage of the landscape-scale carrying capacity, (c) number of boxwood patches disappearing as a percentage of the initial number of boxwood patches, (d) landscape-scale moth biomass, (e) number of patches invaded as a percentage of the number of boxwood patches present, (f) time of moth persistence. The ecologically realistic parameter values are *f* = 120 and *S_m,max_* = 0.5. For each couple of parameters 50 simulations are carried out and results are averaged.

One step further, hypothesis 1b postulates that long-term coexistence is due to asynchronous dynamics, and that moth extinction is due to the synchronisation of the local dynamics. If we artificially ensure that the invasion begins with a moth in each cell, we observe that all boxwood stands are defoliated simultaneously and that the moth disappears globally in a dynamic of type pulse outbreak [Berryman, 1987]: perfect synchronization indeed leads to moth extinction. But if all stands are initially invaded except a single one, this is enough for the occurrence of desynchronisation, and the whole system becomes viable. The moth can therefore disappear due to perfect synchronization, with 100% of the patches invaded simultaneously. However, any other mechanism that globally reduces drastically the resources may also cause moth extinction. Indeed, with high moth fecundity and survival rates (upper-right corners of Figure 2a and f) the moth depletes the resource until its own extinction. This negative effect of fecundity and survival of the BTM was expected from the sensitivity analysis (see *f* and *S_max_* in SI Figure S.9). In contrast to the results obtained with *H. spinipennis–A. dieffenbachii* system, our results indicate that global resource depletion is responsible for moth extinction, rather than synchronisation of local dynamics [Johst and Schöps, 2003]. This is in line with the individual-based model of Uchmański [2019], who found that forest insect pests may go extinct when adult fecundity or larvae survival increase. Within the coexistence regime, the average density of moths and the average intensity of the invasion are insensitive to moth fecundity and survival; the moth either persists at the coexistence density or goes extinct (Figure 2).

Interestingly, the moth population can persist even when it periodically invades 99% of the patches (SI Figure S.8 top), provided that leaves grow back fast enough to prevent global resource depletion. In this case the moth invades a significant proportion of the patches even during troughs (about 12%); the cyclic dynamics are therefore getting closer to a permanent outbreak [Berryman, 1987]. The same process occurred in the model of Uchmański [2019], where leaf growth rate was expressed by a parameter defining the number of years needed for their regeneration. In his model, when the leaf growth rate increased the dynamics of the cyclic outbreak was accelerated, with shorter periods between peaks and troughs. The system then transited to a permanent outbreak for very rapid leaf growth rates. On the contrary, a slow leaf growth rate led to the extinction of the insects (SI Figure S.8). In our model, we also show the positive effect of the wood survival and the leaf productivity on the moth persistence (see SI Figure S.5 and Figure S.9)

#### Spatial structure generates periodical invasions (Hyp 2)

We further postulate that, despite local cycles of extinction and recolonization, the coexistence regime is stationary at the landscape scale (hypothesis 2), a phenomenon called statistical stability [De Roos et al., 1991, Holyoak et al., 2005, Amarasekare, 2008]. Instead, we find that the coexistence regime is periodic at the landscape scale (Figure 3a). A similar pattern has been observed in the *H. spinipennis–A. dieffenbachii* system [Johst and Schöps, 2003] and in Uchmański [2019]. The global period ranges between 20 generations (high leaf growth rate, SI Figure S.8 top) and 40 generations (low leaf growth rate, SI Figure S.8 bottom). In contrast, the invasion of a local patch lasts 5 generations when 1000 moths are introduced at once, and 7 generations when a single moth colonizes the patch (SI Figure S.5). This discrepancy between the local and global timescales suggests that periodicity at the global scale results from the combination of the local time scale and the pace of dispersal. During peaks, between 60 and 99% of the boxwood patches are simultaneously invaded by the moth, and 1-5% during periods of minimal abundance, depending on parameter values. Periodic travelling waves explain this pattern, which have been described in prey-predator systems [Lambin et al., 1998, Sherratt, 2001]. However, such prey-predator systems are locally periodic as well, and long-term coexistence does not require spatial structure. In contrast, in our study system, the periodic waves emerge from the spatial structure, instead of being a mere consequence of local periodicity.

**Fig. 3.**
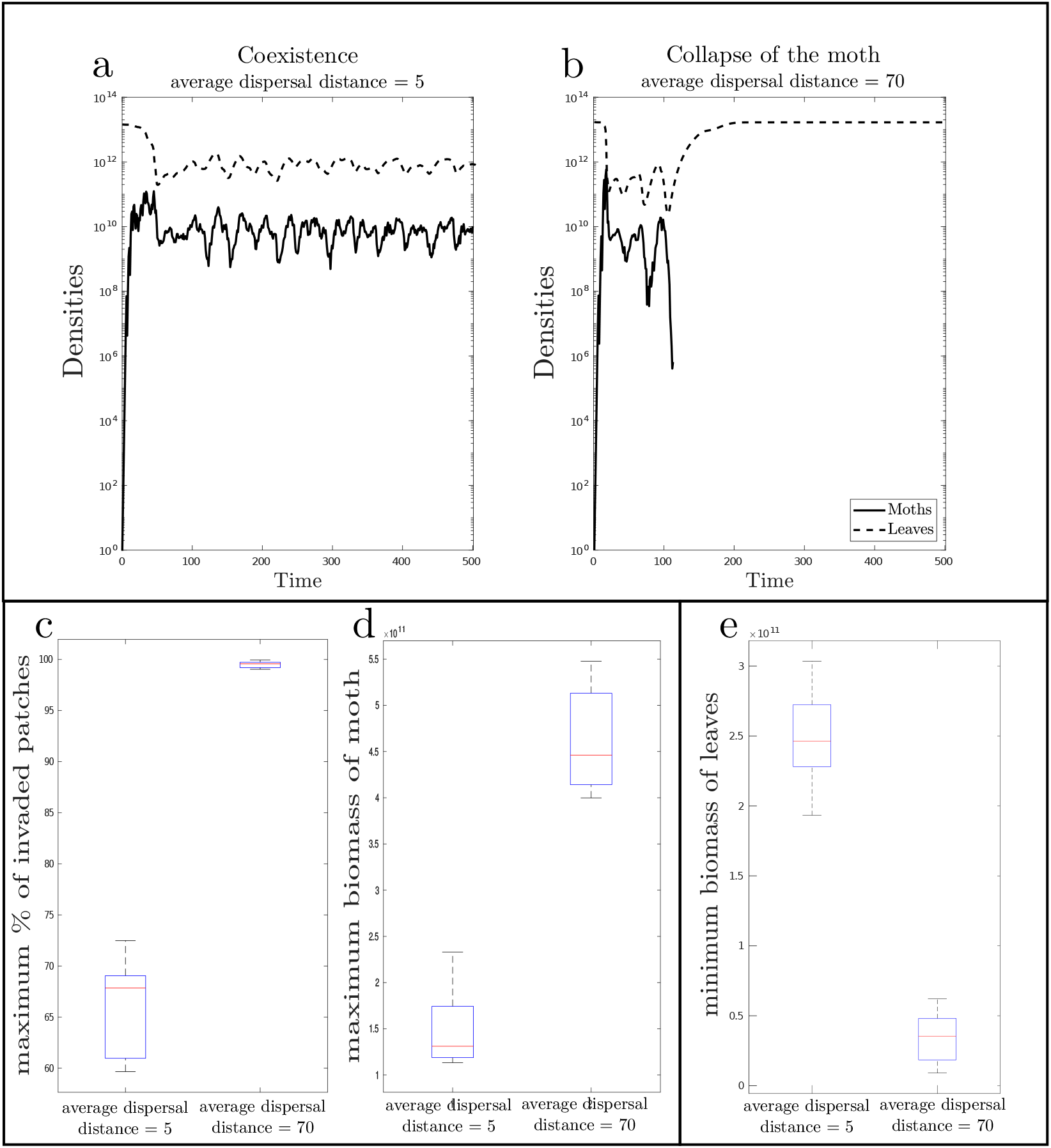
Example of global population dynamics in the case of coexistence (a) and moth collapse (b). (c) maximum % of invaded patches. (d) maximum moth biomass. (e) minimum leaf biomass. Parameters values as in Table 1, except for *r*_0_=2 * 10 ^3^ and for the average dispersal distance *α_d_* which either equals 5 cells (moth persistence) or 70 cells (moth collapse). Boxplots are produced from 50 simulations.

If the mean dispersal distance is very low (1 cell on average, keeping rare long-distance events), the amplitude of the oscillations is also very low (Figure 4b) and the system tends to be statistically stable.

**Fig. 4.**
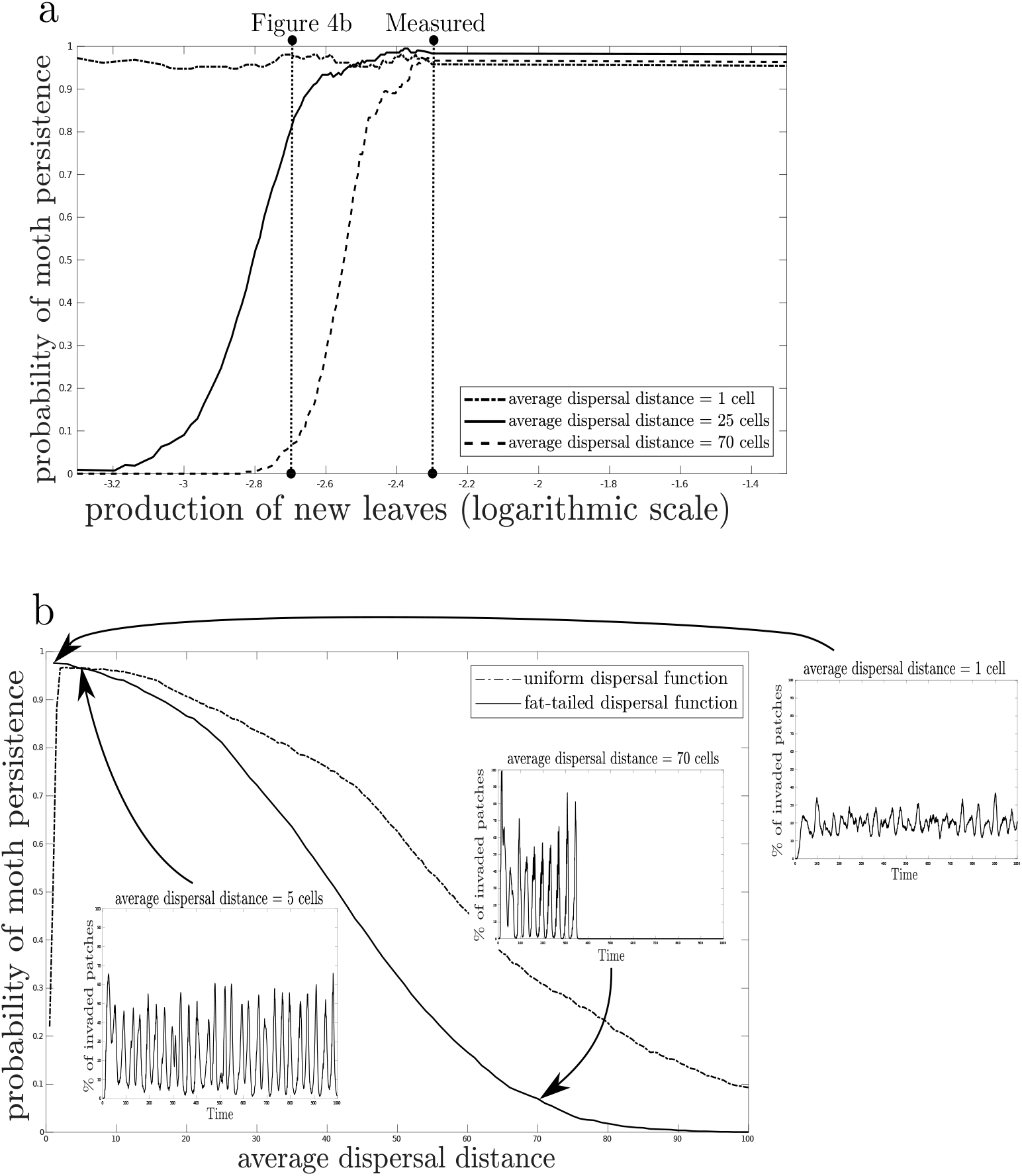
(a) Effect of the rate of new leaves production by the wood (*r*_0_) on the probability of moth persistence. For each tested value, the probability of persistence is obtained by 50 simulations conduct on the realistic landscape with random initial patch of invasion. Three average dispersal distances are tested, a realistic distance of 25 cells, a very short distance of one cell, and a very large distance of 70 cells. (b) Effect of the average dispersal distance on the probability of moth persistence. For each tested value, the probability of persistence is obtained by 50 simulations conduct on the realistic landscape with random initial patch of invasion. Each time two dispersal functions are tested: a fat-tailed function, and an uniform function. The minimum dispersal distance is one cell. The inserts show the percentage of patch invaded over time for three selected average dispersal distances of 1, 5 and 70 cells with the fat tail dispersal function. For each parameter values 50 simulations are carried out and results are averaged.

#### Intermediate intensity of long-distance dispersal prevents BTM from extinction

Hypothesis 3 posits that asynchronous local dynamics require intermediate dispersal distance [Myers, 1976, Myers and Campbell, 1976], because very limited dispersal might prevent suitable boxwood patches from colonization (3a), whereas very long-distance dispersal may synchronize local dynamics (3b). The sensitivity analysis reveals that the moth persistence significantly decreases with an increase of migration parameters, i.e. *M_w,max_* and *α_d_*, migration rate including mortality during dispersal and mean dispersal distance (SI Figure S.9). However, with the measured parameter values we find that the long-term probability of moth persistence is insensitive to dispersal distance. This is shown in Figure 4a, where the measured value for the production of new leaves from wood after total defoliation *r*_0_ equals 5 * 10^−3^ (−2.3 on a log scale). In that case, the moths persist whatever the mean dispersal distance is (1, 25 or 70 cells), because boxwood produces new shoots after total defoliation fast enough to enable moth persistence even when the global leaf biomass is low. Meanwhile, previously infected areas produce enough leaves to support another moth outbreak. Conversely, when *r*_0_ equals 2 * 10^−3^ (−2.7 on a log scale, which may happen under climatic stress for instance), it turns out that the moth does not persist when the mean dispersal distance equals 70 cells (Figure 3b). The moth initially invades 99% of the patches (Figure 3c) and reaches very high densities (Figure 3d). This reduces the global leaf biomass (Figure 3e) and the moth eventually collapses due to resource depletion. In the case of coexistence (Figure 3a) the maximal % of invaded patches is around 60-70%, the maximal moth density is 3 times lower and the minimal leaf biomass is 5 times higher than in the extinction case.

We also find that when the mean dispersal distance is very short (1 cell on average) the probability of moth persistence is not affected by the offshoot production rate. Because of short-distance dispersal, the moth does not generate periods of global invasion and intact patches are always present at the border of its slow moving invasion front. In that case, relatively high offshoot production rates are unnecessary as the moth populations do not rely on the new growth of recently defoliated patches. In contrast, when the average dispersal distance is larger (e.g. 25 and 70 cells), the probability of persistence increases with the offshoot production rate (Figure 4a). In such cases, the moth needs to recolonize recently defoliated patches because during peaks a large proportion of patches are defoliated at the same time. Therefore, relatively high offshoot production rates are necessary to avoid global over-exploitation.

The influence of the mean dispersal distance on moth persistence is therefore studied with a relatively low offshoot production rate, *r*_0_ = 2 * 10^−3^. As expected by hypothesis 3b, frequent long-distance dispersal events lead to the extinction of the moth due to global resource depletion (Figure 4b, continuous line). This effect of long-distance dispersal was not present in Uchmański [2019], this can be explained to the use of a thin-tailed Gaussian kernel characterized by very rare long-distance dispersal events, thus insect pests could not reach the distant trees which had time to regenerate after their last defoliation.

Furthermore, hypothesis 3a posits that the moth goes extinct in the case of limited dispersal, because it would not be able to escape over-exploited patches. However, the fat-tailed dispersal kernel prevents such phenomenon: even when the mean dispersal distance is very low (1 cell), the moth can persist thanks to frequent long-distance dispersal events. Things change when we constrain dispersal to a uniform distribution, which ranges between 1 cell and a given maximum of cells. In that case, when the maximum number of cells is low enough (up to 4 cells), the moth invasion can get stuck in a landscape dead end and it disappears because of local resource depletion, and not global resource depletion (Figure 4b, dotted line). Prey-predator systems subject to limit cycles were also stabilized by limited dispersal in the individual-based model of Cuddington and Yodzis [2000], where limited dispersal reduced the average predation rate and thereby avoided local instability. However, if dispersal is too limited the consumer can go extinct because of a drastic reduction in the rate of predation, as is the case in our model when long distance dispersal events are extremely rare due to uniform dispersal kernel (Figure 4b, dotted line).

The results obtained using the fat-tailed dispersal seems the most plausible, indeed rare long-distance dispersal events mediated either by wind or by human dispersal likely occur in the case of the box tree moth. With a fat-tailed dispersal kernel, multiple invasion fronts are created far away from invaded patches because long-distance dispersal events are frequent [Shaw, 1995]. This created a fragmented landscape of defoliated and intact boxwood, in which the moth does not end up in a dead end. Such a fragmented invasion front can be observed in Europe. The moth was first observed in Germany in 2006/2007; in the same year it spread to Switzerland and the Netherlands and reach France [Feldtrauer et al., 2009] and the United Kingdom [Salisbury et al., 2012] in 2008, Austria in 2009, Italy [Bella, 2013] in 2010, and Portugal, Iran and Armenia in 2016 [Bras et al., 2019]. Long-distance dispersal might be due to the boxwood trade between European countries, and probably to a much lesser extent by the natural dispersal of moths. We therefore expect that in practise only frequent long-distance dispersal can lead to the extinction of the moth, due to global resource depletion.

However, this prediction might be mitigated by a dispersal cost resulting from higher mortality encountered during a long-distance flight, a fertility decline with flight distance or even delayed mating effect that reduces the fitness of moth dispersing at long-distance. Indeed, those mechanisms tends to reduce the high invasion potential of the long-distance dispersal events by limiting the effective dispersal distance. And our model already shows that moth persistence is higher with intermediate-distance dispersal. Moreover, these ecological processes are unlikely to reduce dispersal to the extreme case of a uniform distribution at short distances, and therefore do not significantly affect the persistence of the box tree moth. Indeed, we show that a reduction of the swarm size with dispersal distance gives qualitatively unchanged results on the moth persistence (SI Figure S.4)

#### Fragmented landscape with low amount of resource prevents extinction (Hyp 4)

Next, we predict that the coexistence regime depends on the landscape characteristics, in particular its size (4a) and the proportion of suitable boxwood patches (4b). To do so, we use wrapped landscapes (with no edge effect) of different sizes and different proportions of randomly distributed suitable patches (Figure 5a). Below 200*200 cells, which corresponds to about 12 000 km^2^, coexistence does not occur within the realistic range of parameter values, because previously defoliated patches are quickly recolonized by the moth and lack time to grow back. This induces a global resource collapse and drives the moth to extinction. Above the 200*200 threshold, the larger the landscape is, the more likely the coexistence occurs, in line with hypothesis 4a. In the previous sections, we restrict our model to the French Alps, but we expect that long term coexistence is even more likely on the European scale because larger landscapes favor coexistence. The influence of landscape size is also apparent in the real landscape, with a uniform and short-range dispersal kernel: when the invasion begins in a relatively isolated area of the landscape, coexistence is impaired by small size.

**Fig. 5.**
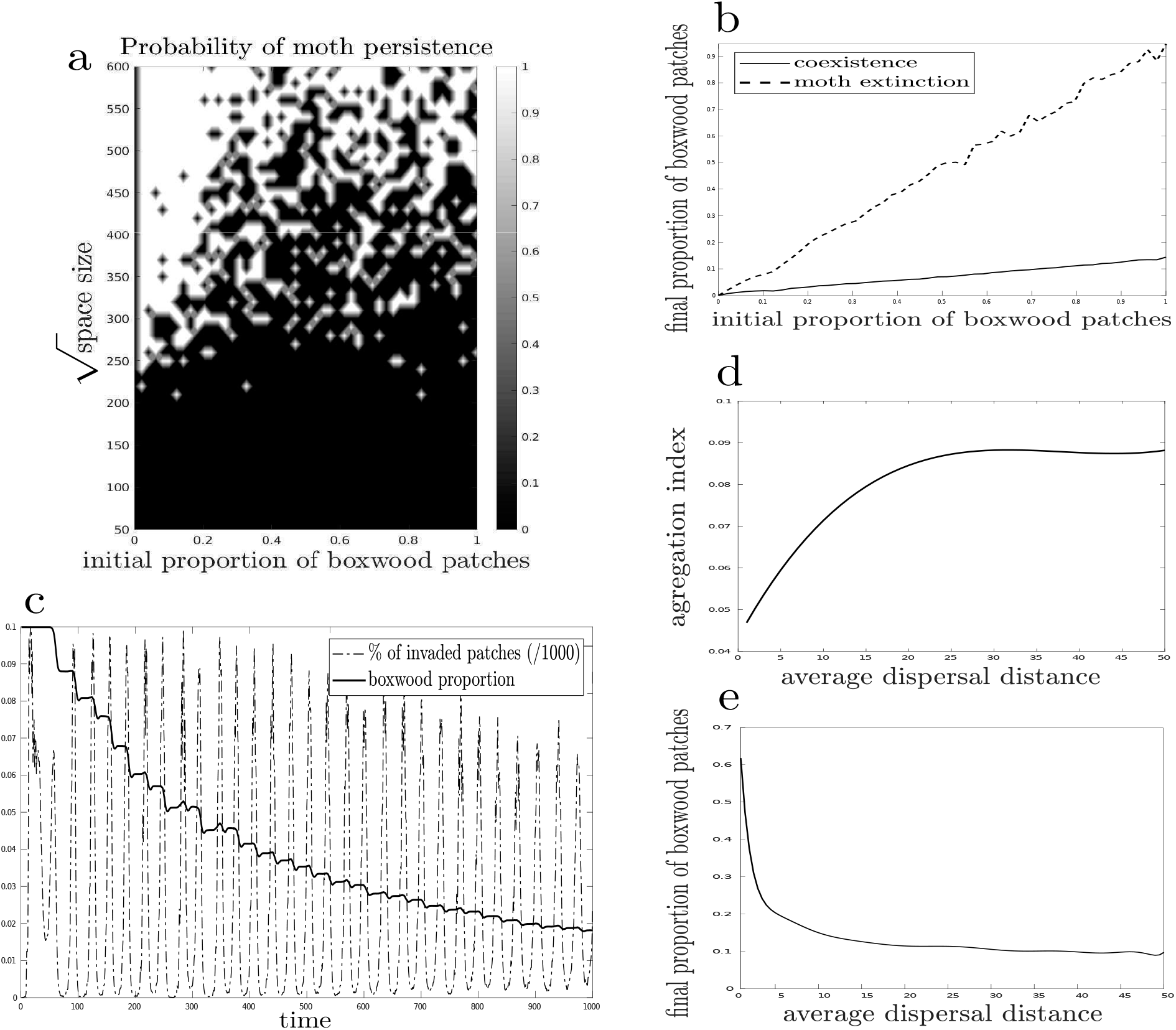
Simulation in theoretical landscapes. (a) The probability of moth persistence depends on space size and on the initial proportion of boxwood patches. (b) In the case of coexistence, the final proportion of boxwood patches is much lower than the initial proportion because moth outbreaks cause patch extinctions. (c) The boxwood proportion in the landscape declines during each moth outbreak. (d) The aggregation index increases along with the average dispersal distance. (e) In coexistence, increasing the dispersal distance reduces the final proportion of boxwood patches in the landscape. All other parameter values are set to the realistic values. For panels a, b, d and e, results are averaged from 50 simulations.

A priori, the effect of the initial proportion of boxwood patches on coexistence is unclear (4b) because on the one hand a higher proportion of suitable patches provides more resources to the moths, while on the other hand it may trigger moth outbreaks which ultimately leads to resource depletion. We find that the latter mechanism is on the driver’s seat: reducing the proportion of boxwood patches increases the probability of moth persistence (Figure 5a). More precisely, the landscapes larger than 400*400 cells filled with less than 20% of boxwood patches almost always allow coexistence. In such cases, most dispersal events fail to find suitable patches, lowering the moth population density, which in turn leaves more time for leaves to grow back. On the contrary, a high proportion of suitable patches results in high moth densities which leads to global over-exploitation, despite a potentially higher leaf biomass. Thus, we postulate that any environmental changes due to climate changes or human activities, that reduces boxwood biomass would not necessarily drive the BTM towards extinction.

The coexistence regime has interesting consequences on the final proportion of boxwood patches, which corresponds to the initial proportion minus the proportion of complete withering patches due to over-exploitation. Under coexistence, the final proportion increases linearly with the initial proportion with a weak slope of about 0.1, whereas the slope is close to 1 in the case of moth extinction (Figure 5b). This indicates that the local extinction of boxwood patches is not responsible for global moth extinction. Instead, the final proportion of boxwood patches is a long-term consequence of coexistence and results from their gradual death (Figure 5c). During each moth outbreak, a small proportion of the boxwood patches disappears due to over exploitation. Then, right after the outbreak a few boxwood patches are recolonized from neighbouring patches (only previously occupied patches can be recolonized), which induces a clustered distribution of the boxwood patches. As a result of clustering, the aggregation index is always positive, which indicates that the landscapes created by long-term coexistence are more aggregated than random landscapes. Boxwood patches relatively isolated in the initial landscape experience larger extinction rates and create holes in the landscape. In contrast, areas where boxwood patches are initially more abundant persist more often, which creates clusters. Moreover, the aggregation increases with the average moth dispersal distance (Figure 5d). Boxwood patches favor moth outbreaks: increasing the boxwood patches proportion over the landscape produces severe outbreaks. This induces apparent competition between boxwood patches because of their shared pest. With low dispersal distance, apparent competition between patches is mainly local, which limits the formation of clusters. Instead, with high dispersal distance apparent competition is global and the aggregated pattern results from the interplay between local facilitation (recolonization of boxwood patches is purely local) and global competition, as in many spatially self-structured systems [Kéfi et al., 2007, 2008]. This is confirmed by simulations where boxwood recolonization is global, in that case the aggregation process vanishes (details not shown).

We further explore how the aggregation process creates boxwood clusters of different sizes, for various dispersal distance. To do so, we start with landscapes which are initially filled with boxwood patches, and run simulations after invasion by the moth. At the end of the simulations, we fit a power-law model to the final distribution of the cluster sizes (SI Figure S.9 top). Clusters smaller than 5 boxwood patches are excluded from the fit. We find that small dispersal distance leads to a cluster size distribution closer to a power-law than large dispersal distance. When the dispersal distance is large (10 to 50 cells), the cluster size distribution follows a truncated power law (SI Figure S.9 top), which indicates that large clusters are under-represented. Large dispersal distance leads to an increase of herbivory, which produces two distinct effects. On the one hand, it favors aggregation due to global apparent competition, as discussed earlier (Figure 5d). On the other hand, it increases the death rate of boxwood patches and thus reduces the final proportion of patches in the landscape (Figure 5e). This is why under large dispersal distance (10 to 50 cells) the final landscapes has a homogeneous aspect of equally spaced small clusters (SI Figure S.9 bottom).

These regular patterns are similar to Turing instabilities [Turing, 1990, Murray, 2001] and result from “scale-dependent feedbacks” which combine short-distance positive feedbacks and long-distance negative feedbacks [Rietkerk and van de Koppel, 2008]. In the present case, short-distance positive feedback correspond to local facilitation of boxwood due to recruitment while apparent competition between boxwood stands because of their shared pest mirror long-distance negative feedbacks. Several studies have investigated how spatial patterns can emerge from such scale-dependent feedbacks in a variety of ecological scenarios, such as plant-water interactions [Klausmeier, 1999, von Hardenberg et al., 2001, Rietkerk et al., 2002, Meron et al., 2004, Kéfi et al., 2010], plant-plant interactions [Lejeune et al., 1999], or predator-prey interactions [Levin and Segel, 1976, Solé and Bascompte, 2012]. It has been shown that such spatial patterns emerge in predator-prey systems when the predator has a larger dispersal capacity than the prey [Gurney et al., 1998, de Roos et al., 1998]. We demonstrate here that this can also be the case in the context of a plant-herbivore system, using a model calibrated empirically.

#### Implications for management

First, the most important finding for management purposes is that the moth heavily impacts boxwood stands. With the estimated parameter values, we find that in the study area only 10% of the initial boxwood biomass remains after invasion (Figure 2b) and that 50% of the original boxwood patches completely disappear (Figure 2c), which represents 2414 square kilometres in the French Alps. However, these quantitative predictions should be taken with caution. In particular the sensitivity analysis (SI Figure S.9) shows that the characteristics of boxwood are determining factors in the model, so a better understanding of the biology of boxwood and its response to defoliation seems essential to gain further insights into the impact of the box tree moth invasion. Under low moth fecundity and high caterpillar survival, the moths could persist longer in heavily defoliated patches. The severe decrease in box tree biomass can impact the many species associated with boxwood, as well as the ecosystem services provided by the shrub (i.e sediment trapping and water storage) [Mitchell et al., 2018]. In stands where boxwood is severely weakened or extinguished, recolonization by neighbouring patches may be prevented by pioneer plants, potentially other invasive species such as *Buddleja davidii* or *Ailanthus altissima*, in a kind of ‘invasion meltdown’ process [Simberloff and Holle, 1999].

Next, the periodic invasion dynamics can lead to confusion regarding the persistence of the box tree moth. A period of low overall abundance should not be confused with a decrease in invasion, and moth control methods should take periodic invasion dynamics into account. Remote sensing methods may be appropriate in order to detect the few boxwood stands that provide refuge under low moth abundance [Kerr and Ostrovsky, 2003]. We suggest that detecting stands of undefoliated boxwood that allow moth persistence during a period of low abundance could provide an interesting management strategy, since control efforts could be increased on these particular patches.

Finally, management actions might consider preventing anthropogenic long-distance dispersal. However, we find that only very limited dispersal could lead to moth extinction, which occurs with a uniform distribution of dispersal events of no more than 4 cells (Figure 4b, dotted line). As soon as dispersal is higher, the moth could escape dead ends in the landscape and therefore persists. Even if anthropogenic long-distance dispersal is prevented, a few natural long-distance dispersal events might ensure moth persistence. It is therefore unlikely that management actions limiting dispersal can be able to eradicate the moth. However, such actions can reduce the impact on boxwood stands, since we find that long-distance dispersal increases the extinction rate of boxwood patches (Figure 5e).

Three scenarios may occur after the invasion of box tree moth in Europe: extinction of both species, extinction of the moth only, or coexistence. Our theoretical approach combined with field and experimental data suggests that coexistence is the most likely outcome, with cycles of moth outbreaks and crashes. Coexistence comes along with a severe reduction of boxwood biomass at the landscape scale: boxwood stands may therefore become closer to brown than green. Moth extinction can also occur, which indicates that the invasion dynamics of exotic pests can be mitigated even in the absence of predators and effective plant defenses.

We further show that plant-herbivore coexistence through spatial effects does not require poorly mobile wingless species, as in our model a wide range of dispersal values result in coexistence. Coexistence occurs in large landscapes, long-distance dispersal thus requires a spatial scaling-up for the persistence of the system. Particularly intense long-distance dispersal nevertheless leads to herbivore extinction, provided that plant grows back slowly. In that case, the herbivore depletes its resources at the global scale, which leads to its own extinction even without a complete synchronization of the local dynamics. Finally, coexistence is easier in patchy landscapes because unsuitable patches increase moth mortality during dispersal and thereby reduce the global insect population size. Interestingly, when plants disperse locally the spatial dynamics of the system lead to the formation of such a patchy landscape, with relatively small plant patches evenly distributed in the landscape. The system thus creates its own stability conditions.

## Author contributions

CG, SI and JG originally formulated the idea, CG, SI, JG and LL developed methodology, LL conducted fieldwork, SI, JG and LL developed the mathematical model, LL performed the numerical analyses, all authors participated in writing the manuscript.

## Acknowledgements

- We thank all the student interns who participated in both field and experimental work: Aristide Chauveau, Alison Dilien, Jessica Barbe and Océane Guillot. We also thank Elisabeth Tabone and Thomas Defferier (INRAE) for their mass measurements on their farmed caterpillars, and Audrey Bras (INRAE) for the flight distance data and fruitful exchanges.
- This article is financially supported by AAP USMB and FREE federation. The PhD scholarship of L. Ledru is funded by the French Ministry for Education and Research. J. Garnier acknowledges GLOBNETS project (ANR-16-CE02-0009).

