## Supplemental Materials for "Spatial structure of natural boxwood and the invasive box tree moth can promote coexistence"

#### 1 Model description

**Reproduction.** The wood growth function $F_w$ is constructed using a Ricker model whose intrinsic growth rate $r_w(\rho_n)$ is defined as the balance between the production of new wood $b_w$, which critically depends on the density of leaves per unit of wood $\rho_n$, and the mortality induced by severe defoliation $d_w$.

$$F_w(w,\rho) = \begin{cases} \exp(r_w(\rho)(1-\frac{w}{W_{max}})) & \text{if } r_w(\rho) > 0 \\ \exp(r_w(\rho)) & \text{if } r_w(\rho) \leq 0 \end{cases} \qquad \text{with} \quad r_w(\rho) = b_w(\rho) - d_w(\rho)$$

The production function $b_w$ and the mortality function $d_w$ takes the following form :

$$b_w(\rho) = \frac{\rho^{\beta_r}}{\rho^{\beta_r} + \theta_r^{\beta_r}} r_{w,max} \qquad\qquad d_w(\rho) = (1 - d^{\frac{1}{\rho}}) r_{w,min} \tag{1}$$

where $\beta_r$, $\theta_r$ and $d$ are shape parameters.

**Survival.** The survival function of the leaves is defined by

$$S_l(\mu) = v\,(\sigma_l^{\mu}) \tag{2}$$

where $\sigma_l$ is a shape parameter.

The wood mortality due to consumption increases with both the BTM pressure, $\mu$ and the BT health $\rho$. The wood mortality saturates to $d_{max}$ when the foliage is abundant (large ratio $\rho \geq 1/3$). However, when the foliage is small (low ratio $\rho \leq 1/3$), the bark consumption occurs while the superficial wood is available. Thus recently defoliated boxwood with small bark coverage ($\rho$ close to 0) cannot be consumed by the larvae.

$$S_w(\mu, \rho) = 1 - \frac{\mu^{\beta_s}}{\mu^{\beta_s} + \theta_s^{\beta_s}} D_{max}(\rho) \tag{3}$$

where $\beta_s$ and $\theta_s$ are shape parameters and $D_{max}$ is the maximal mortality rate which grows linearly with $\rho$ until a threshold 1/3, at which point it saturates to its critical value $d_{max}$.

$$D_{max}(\rho) = \begin{cases} 3d_{max}\,\rho & \text{if } \rho < 1/3 \\ d_{max} & \text{if } \rho \geq 1/3 \end{cases} \tag{4}$$

This step function takes into account that the consumption of superficial wood depends on the presence of available softwood, and thus a certain amount of foliage. The threshold value 1/3 for BTM pressure $\rho$ corresponds to the approximate ratio when there is as much foliage as wood, the density of the wood being three times greater.

Survival of BTM during the larval stage depends mainly on the amount of available resource per larva $\mu$. If the larva has enough available resource to complete its six stages, it will evolve into a moth, while a lack of resource during its growth will cause its death. The survival rate also takes into account intraspecific competition for resource caused by interference between the larvae. The survival function is defined using a shape parameter $\sigma_m$ as

$$S_m(\mu) = \begin{cases} S_{m,max}\left(1 - (\sigma_m)^{\frac{1}{\mu}}\right) & \text{if } \mu < 2 \\ 0 & \text{if } \mu > 2 \end{cases} \tag{5}$$

The threshold $\mu = 2$ fits a given situation that occurs when there is a shortage of resource for the larvae, even if some larvae die during their development.

**Dispersal phase of BTM** The migration rate $M_m$ of BTM depending on $\mu$ takes the following form :

$$M_m(\mu) = (1 - \delta^{\mu}) M_{m,max} \tag{6}$$

where the maximal dispersal rate $M_{m,max}$ takes into account mortality during the dispersal. Thus the number of dispersal events at each location is given by $\frac{M_m(\mu)m}{1000}$.

**Dispersal of BT.** The dispersal rate of BT depending on the foliage ratio per BT ($\rho$) takes the following form :

$$M_w(\rho) = \begin{cases} (\omega_w)^{1/\rho} M_{w,max} & \text{if } \rho > 1/3 \\ 0 & \text{otherwise} \end{cases} \tag{7}$$

### 2 Figures

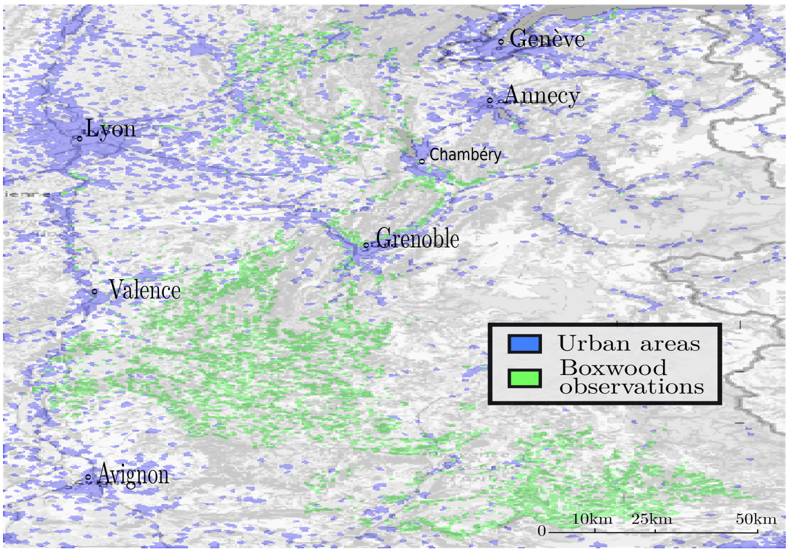


Figure S.1: Simulation space of the cellular automaton representing the distribution of boxwood in the French alpine arc, based on data from the National Alpine Botanical Conservatory. 570 by 351 cells of 29 hectares each, about 58 000km$^2$.

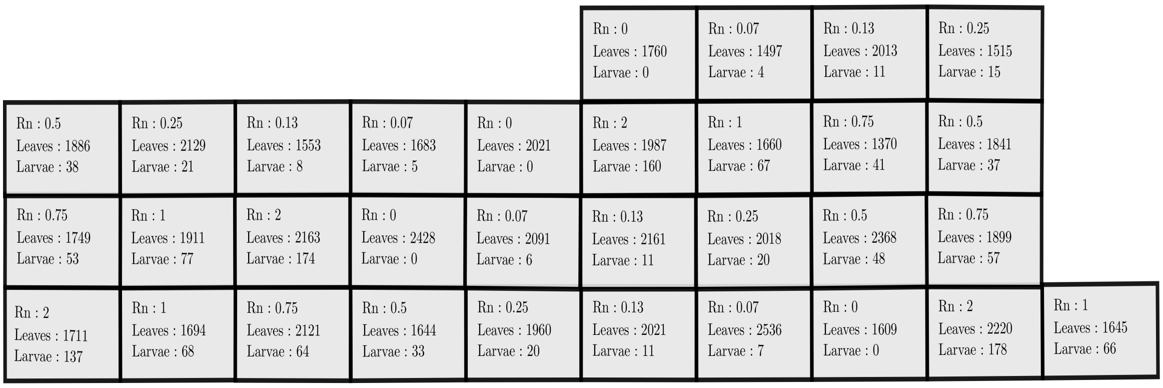


Figure S.2: Schema of the mesocosm manipulation. $R_n$ represents the competition for the resource. The spatial layout is due to the topography of the installation site

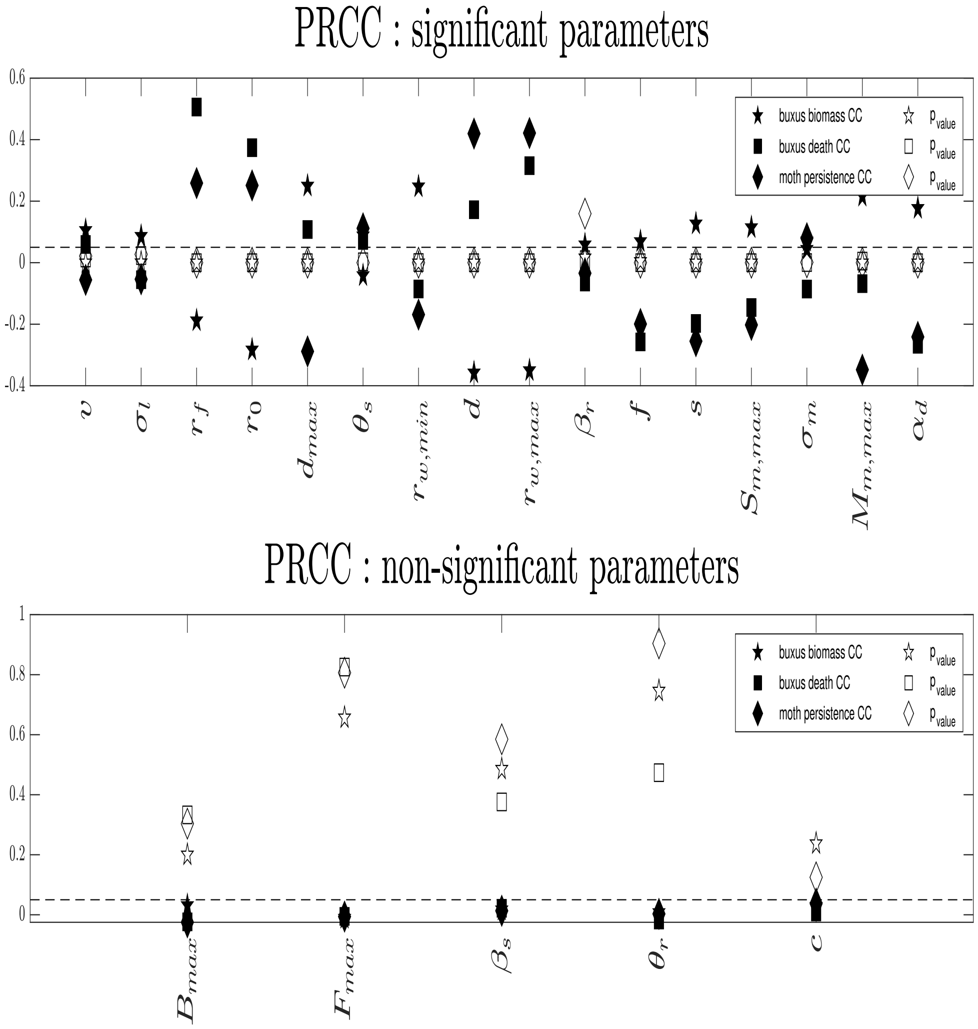


Figure S.3: Partial Rank Correlation Coefficient analysis. The analysis is conducted on 21 parameters and three proxies: boxwood biomass as a percentage of the carrying capacity, percentage of dead boxwood patches and probability of box tree moth. A parameter is considered significant as soon as it has a significant p-value for a proxy. The Correlation Coefficient (CC) shows the relationship between the paramter and each proxy.

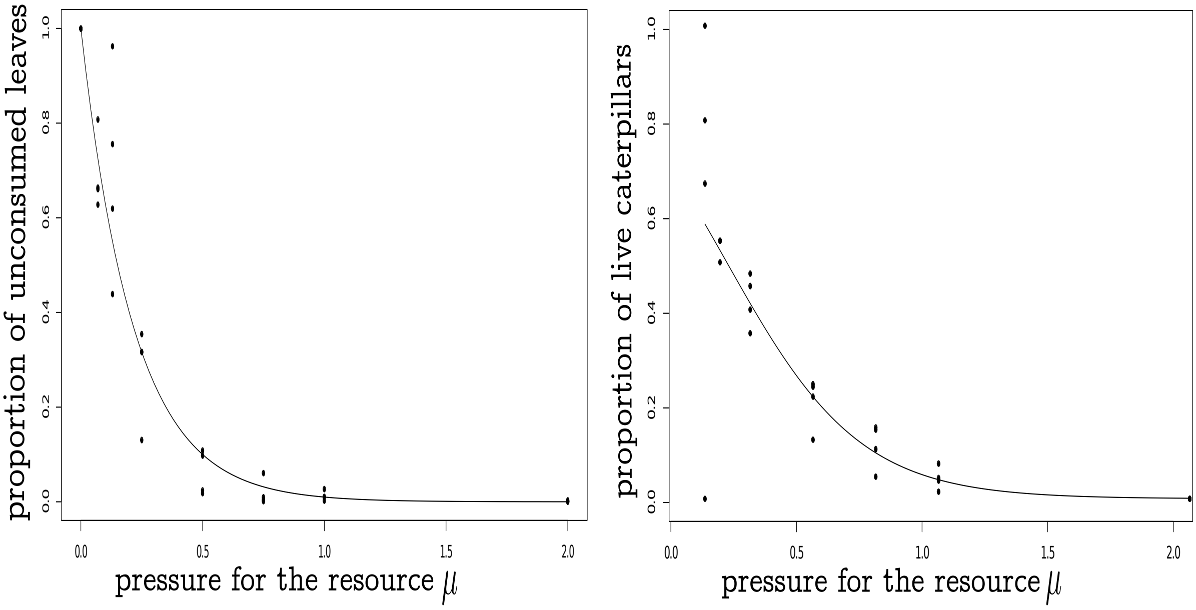


Figure S.4: Main results of mesocosm manipulation with measurement of defoliation intensity and box tree moth survival according to competition for the resource $\mu$. (a) leaf consumption function $\frac{S_l(\mu)}{v}$. (b) box tree moth survival function $S_m(\mu)$.

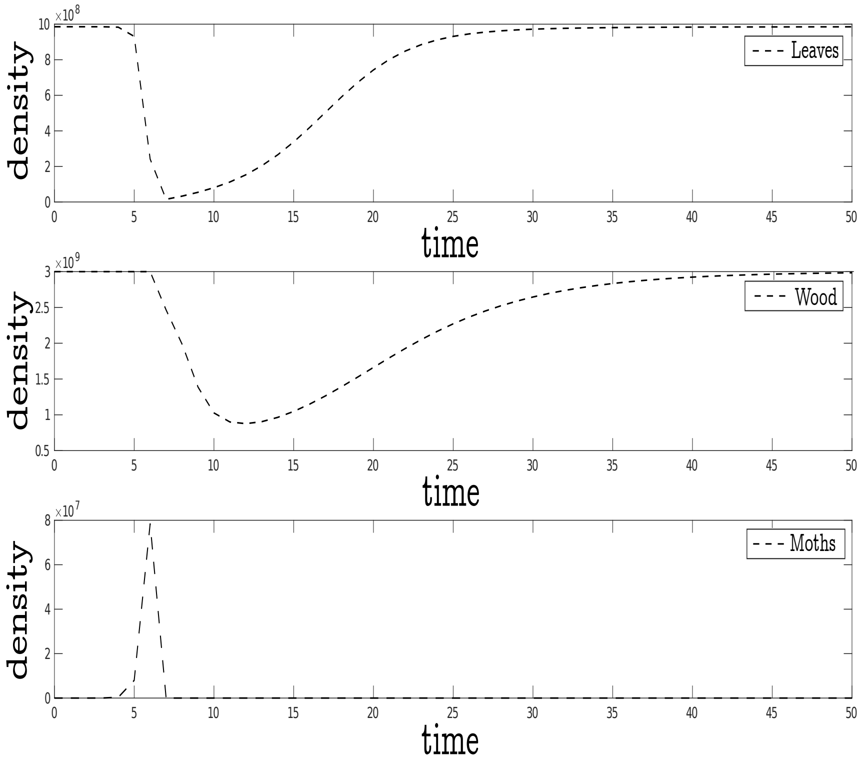


Figure S.5: Visualization of invasion dynamics in the local model. Simulation with ecologically realistic parameters given in Table 1

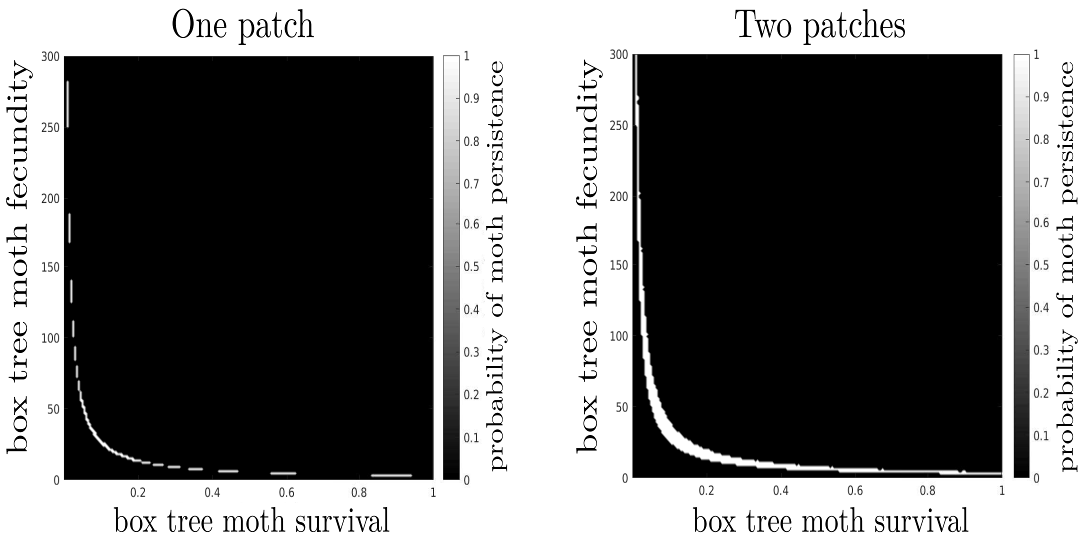


Figure S.6: Final state map for the mean field model (left) and the double-patches model (right) as a function of fertility $f$ and maximum survival $S_{m,max}$. The parameter values considered ecologically real are $f = 120$ and $S_{m,max} = 0.5$.

In the two-patch case, patches are initially saturated with boxwood, and the patch 1 contains just one moth. This spatialization of the model into two patches leads to new peculiar and rare end states at the frontier between coexistence and extinction which are not represented. We observe five states: i) overall disappearance of the moth and persistence of the boxwood (black) ii) global coexistence of the two species (white) iii) extinction of the moth and the boxwood in patch 1 and coexistence in patch 2, iv) extinction of only the moth in patch 1 and coexistence in patch 2, v) coexistence of the two species in patch 1 and the absence of moth in patch 2. It is important to note that cases iii) and iv) are not actually final states; the increase in simulation time shows that in fact the moth population in patch 2 decreases slowly until its extinction. For iii), after the extinction of the moth, the boxwood recovers in patch 2 until it reaches a state where it could disperse and recolonize patch 1. Moreover, before the moth extinction, during its decline the dispersal to patch 1 is still active, though low. Thus, in this situation patch 2 is a source of moths, and patch 1 is a well since migrating moths find no resource and die immediately. On the other hand, for iv), moth densities are too low in patch 2 and there is no dispersal towards patch 1. Cases iii) and iv) are not asymptotic states, but they are still taken into account because on an ecologically reasonable time scale they are actually present.

Nevertheless, except for the states i), global extinction of the moth, and ii), global coexistence of the two species, the other states are highly anecdotal because they correspond to only a few very precise parameter values and are almost invisible in the parameter space. What is remarkable in the space of parameters, however, is the slight expansion of the space corresponding to the coexistence of boxwood and moth

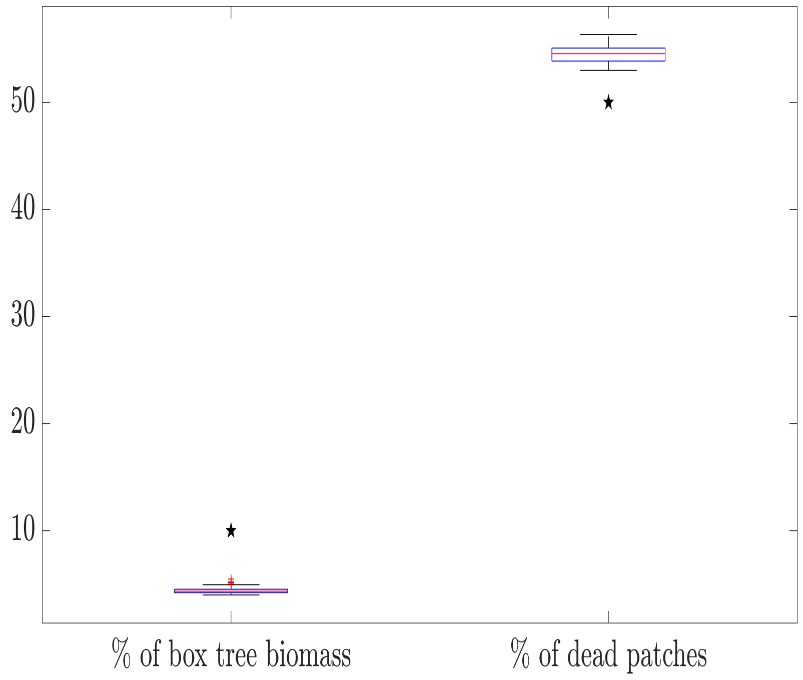


Figure S.7: Comparison of simulations including additional ecological processes (boxplots), i.e. possible effects of elevation and dispersal distance, with reference simulations (stars). Elevation is assumed to reduce egg survival (thus fecundity), larvae survival and the nutritional quality of boxwood, up to a maximum reduction of 50%. One thousand simulations are performed on real space, each time a uniform random matrix is drawn to determine the elevation of each cell (with ten replicates). We measure as proxies the persistent boxwood biomass as a percentage of the carrying capacity and the percentage of bead boxwood patches. In addition, the size of the dispersal swarms reduces linearly with the dispersal distance to a minimum of one individual.

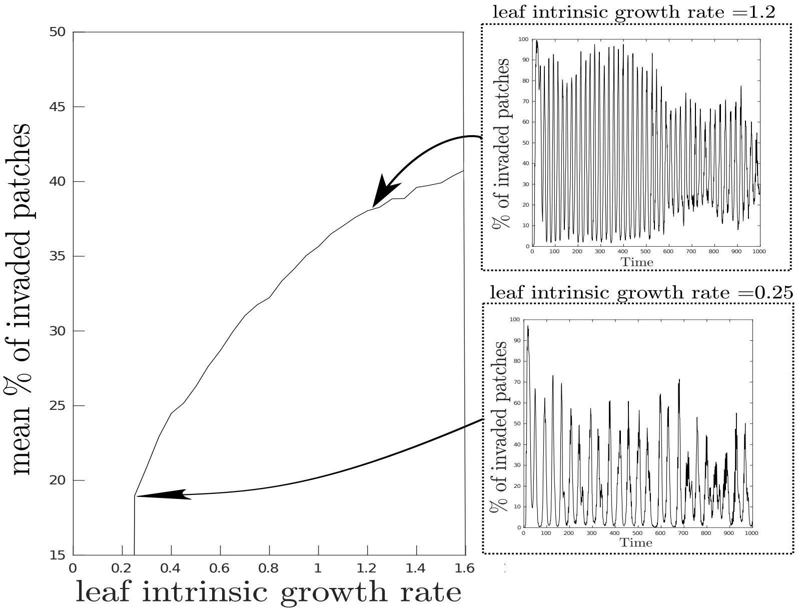


Figure S.8: Effect of the leaf intrinsic growth rate on the mean percentage of invaded patches. The percentage drop to zero when the moth do not persist in these conditions (when $r_f < 0.22$). The two inserts show the invasion dynamics for two growth rate values. For each value of leaf intrinsic growth rate 50 simulations are carried out and results are averaged.

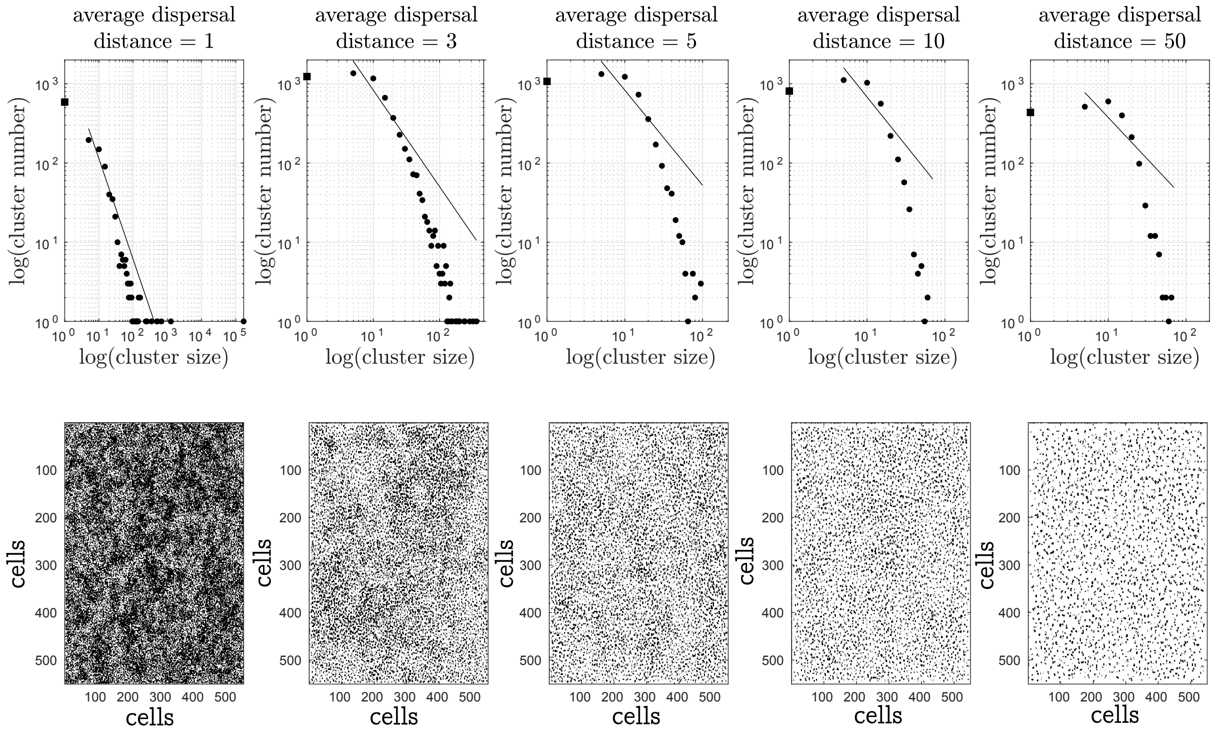


Figure S.9: Effect of moth dispersal distance (increasing from left to right) on the size distribution of boxwood clusters at the end of the simulations (5000 time steps), in large theoretical landscapes of 550*550 cells. Top: regression lines correspond to the fit to a power-law distribution. The first size class (black square) has been excluded from the regression. Bottom : snapshot of the landscape at the end of the simulation, black cells indicate live boxwood patches. All other parameter values are set to the realistic values.
